# Doing nothing and what it looks like… Inactivity in fattening cattle

**DOI:** 10.1101/833012

**Authors:** Sara Hintze, Freija Maulbetsch, Lucy Asher, Christoph Winckler

## Abstract

Animals kept in barren environments often show increased levels of inactivity and first studies indicate that inactive behaviour may reflect boredom or depression-like states. However, inactivity does not necessarily reflect negative welfare and can even be a sign of positive welfare, for example in terms of relaxation. To date, knowledge of how to reliably differentiate between positive and negative states associated with inactivity is scarce and methods to identify different forms of inactivity are thus warranted. To this end, we developed an Inactivity Ethogram including detailed information on the postures of different body parts (Standing/Lying, Head, Ears, Eyes, Tail) for fattening cattle, a farm animal category often kept in barren environments. The Inactivity Ethogram was applied to Austrian Fleckvieh heifers from intensive, semi-intensive and pasture-based husbandry systems. Three farms per husbandry system were visited twice; once in the morning and once in the afternoon to cover most of the daylight hours with our observations. During each visit, 16 focal animals were continuously observed for 15 minutes each (96 heifers per husbandry system, 288 in total). Moreover, the focal animals’ groups were video recorded to later determine the inactivity level on the group level. Group level and focal animal data were analysed with (generalised) linear mixed-effect models with husbandry system as fixed effect and (group nested in) farm visit nested in farm as random effects. Husbandry system did not affect group level inactivity or the time the different postures were adopted (with the exception of asymmetrical ears, which were more prevalent in intensive than in semi-intensive than in pasture systems). In addition to the analysis of the time the single postures were observed for, simultaneous occurrences of postures of different body parts (Standing/Lying, Head, Ears and Eyes) were analysed using the machine learning algorithm cspade to provide insight into co-occurring postures of inactivity. Frequently co-occurring postures were generally similar between husbandry systems, but with subtle differences. The most frequently observed combination in intensive and semi-intensive systems was Lying with Head up, Ears backwards and Eyes open whereas in pasture systems it was Lying with Head up, ears low and eyes closed. To conclude, both the Inactivity Ethogram (including the description of detailed postures) and the machine learning algorithm cspade (for identifying frequently co-occurring posture combinations) are promising tools to understand how combinations of postures may be used to distinguish between different affective states associated with inactivity.

## Introduction

Many of the animals we keep on farms, in labs or in zoos live in monotonous and often barren environments. Under conditions that lack changing stimuli, animals often exhibit active behaviour which is indicative of poor welfare, including abnormal repetitive behaviour (e.g. Würbel, 2001) or re-directed behaviour towards conspecifics (e.g. tail-biting in pigs: Scott et al., 2007). Such behaviours are rather striking and have thus been studied intensively across species. In contrast, relatively little attention has been devoted to the high levels of inactivity often described in animals kept in barren environments (Fureix and Meagher, 2015).

Studies of pigs, mice and minks housed in either enriched or barren environments found that under barren conditions, animals spend more time being inactive (Beattie et al., 2000; Bolhuis et al., 2006; Fureix et al., 2016; Haskell et al., 1996; Meagher et al., 2017; Meagher and Mason, 2012). Moreover, pigs and minks experiencing a change from enriched to barren conditions show more inactive behaviour when compared to animals housed in barren conditions throughout the experiment (Bolhuis et al., 2006; Meagher et al., 2013). In veal calves, it has been shown that the feeding regime can also affect the duration the animals are inactive. Calves being fed a barren diet lie more idle than calves receiving an enriched diet, and calves with *ad libitum* access to straw stand less idle than individuals with no additional straw provided (Webb et al., 2017). In sum, these studies indicate that barren conditions lead to increased levels of inactivity, but whether this inactivity is simply the result of “having nothing else to do” or whether it is indicative of more negative states is largely unknown.

Anecdotally, it has been hypothesised that inactive animals are bored (Wood-Gush and Beilharz, 1983) and that being inactive is a “cut-off” strategy of the animal “to isolate itself from an unsuitable environment” (Pearce et al., 1989, p. 35). Only recently, systematic studies on what inactive behaviour really means for the animal have been conducted. From these studies we know that the level of inactivity in mice tends to predict immobility in the Forced Swim Test, a test used to assess depression in rodents (although the validity of this test has recently been questioned (Reardon, 2019)). Moreover, “withdrawn” states of inactivity in horses (as defined in Fureix et al., 2012) are associated with anhedonia (Fureix et al., 2015), one key symptom of depression. While these studies indicate a positive relationship between inactivity and depression-like states, others have not found such an association (Harvey et al., 2019), or have shown a link between inactive behaviour and boredom-like states. In two independent studies, Meagher and colleagues demonstrated that inactivity in mink is positively associated with some measures of exploration of differently valenced stimuli (Meagher et al., 2017; Meagher and Mason, 2012). Increased exploration of all kinds of stimuli (positive, ambiguous and negative) was *a priori* operationally defined as indicating boredom. A similar approach was taken by Webb and colleagues who investigated the reaction of calves receiving different feeding regimes towards novel stimuli, but no relationship between inactivity levels and the calves’ responses in the novel object task was found (Webb et al., 2017).

Taken together, the described studies indicate that increased levels of inactivity in barren environments reflect some kind of negative welfare, but how being inactive is perceived by the animals is still poorly understood. One explanation for this lack of knowledge is probably due to the heterogeneous nature of inactive behaviour. Being inactive can reflect different forms of negative welfare, depending on the context. A study in mink by Meagher and colleagues, for example, indicated that lying in the nest-box may indicate fear or anxiety, while lying awake in the open cage may reflect a boredom-like state (Meagher et al., 2013). Moreover, being inactive does not necessarily reflect negative welfare; in contrast, it can even be a sign of positive welfare, including relaxation, post-consummatory satisfaction and ‘sun-basking’ (as reviewed in Fureix and Meagher, 2015). Different forms of inactive behaviour in different contexts may thus have even opposite meanings with respect to animal welfare, rendering its study very challenging.

A further challenge is that it is difficult to define what being inactive looks like in different species. Most studies define an animal as being inactive if it is standing or lying (or sitting in the case of pigs), often accompanied by the specification that the animal’s eyes must be open (e.g. Beattie et al., 2000; Meagher and Mason, 2012). However, other definitions exist as well. Bolhuis and colleagues, for example, described pigs as being inactive when lying with eyes either open or closed, but differentiated between these two forms (Bolhuis et al., 2006, 2005). Sitting and standing were also recorded in these two studies, but were not included in the definition of inactivity. Webb and colleagues differentiated between lying and standing either actively or idle, the latter being described as “displaying no behaviour with an obvious function” (Webb et al., 2017). To our knowledge, only one study has further specified different forms of being inactive (besides distinguishing between standing and lying) by recording three different lying postures in mink (Meagher et al., 2013). Apart from this, detailed descriptions of inactive animals are lacking.

To fill this gap, we aimed to describe in detail inactivity in fattening cattle, *Bos taurus*, an animal category often kept in barren conditions. Precisely, the aims of our study were 1) to develop an Inactivity Ethogram for fattening cattle encompassing a description of the animals’ basic body postures, lying postures, head and ear postures as well as the closure of their eyes and tail movements, and 2) to apply this Inactivity Ethogram on farms with intensive, semi-intensive and extensive husbandry systems aiming to investigate the time the single postures were adopted and the co-occurrence of the different postures to better understand the complexity of inactive behaviour.

## Animals, material and methods

### 1) Development of an Inactivity Ethogram

Aiming to capture different aspects of body language of inactive cattle, we developed an Inactivity Ethogram. To this end, we first watched existing video clips of fattening cattle to get an impression of the various postures these animals show in order to draft a first ethogram. We then amended our ethogram during four pilot farms visits, during which we observed fattening cattle kept in different husbandry systems ranging from intensive (fully-slatted floor pens) to semi-intensive (feeding and activity area plus straw-bedded lying area) to extensive systems (pasture). Different types of husbandry systems were chosen to encompass a range of different conditions and thus potentially different forms of inactive behaviour. Our pilot visits lasted from the morning to the late afternoon to cover different activity and thus inactivity phases of the animals.

Besides amending the ethogram, we also worked out a definition for being inactive during our pilot visits. An animal was defined as inactive when it fulfilled two requirements. First, it had to either stand or lie. Since an animal can also stand or lie in an active manner (e.g. Harvey et al., 2019; Webb et al., 2017), we specified all movements that classified the standing or lying animal as being either inactive or active (Table 1). For movements that could be classified as both, e.g. scratching with one foot, the general rule was that an animal was recorded as active when this movement occurred more than two consecutive times, while movements that were repeated only once were interpreted as “impulsive” movements, e.g. kicking after flies, and the animal was thus still recorded as being inactive. Second, an animal had to be inactive as defined above for at least 30 seconds before it was classified as being inactive. Both requirements were defined, adapted and their application was trained during our pilot farm visits to ensure consistent data recording during the experiment. In addition to the postures defined for inactive cattle (Table 2), we included some categories of active behaviour in our ethogram to be able to place the inactive behaviour into the context of the shown active behaviour (Table 3).

**Table 1.** Movements classifying standing and lying animals as being inactive or being active

|  | **Movements that classify the animal as being/remaining inactive** | **Movements that interrupt inactivity; the animal is recorded as being active** |
| --- | --- | --- |
| Body movements | • Maximum two steps forward or backward<br>• Single movement of one leg, including a kick after flies<br>• Lying down or standing up without further movements<br>• Stretching while standing or lying<br>• Skin twitching<br>• Urinating or defecating<br>• Any tail movements | • Scratching self with one foot more than twice<br>• Scratching self on barn equipment or objects on pasture<br>• Licking self more than twice<br>• Sniffing an object or conspecific |
| Head movements | • Head shaking (while standing or lying)<br>• Snapping after flies by quickly throwing the head towards one side of the body<br>• Ear movements<br>• Eye blinks<br>• Yawning<br>• Coughing<br>• Sneezing | • Flehming |
| Vocalisations | • Humming | • Mooing |
| Interactions with a conspecific | • Being licked without obvious reaction<br>• Being nibbled without obvious reaction<br>• Being mounted<br>• Receiving a head butt without obvious reaction<br>• Being displaced and being inactive thereafter | • Licking a conspecific<br>• Nibbling on a conspecific<br>• Mounting a conspecific<br>• Head butting<br>• Displacing a conspecific |

Note that some movements could be classified as both the animal still being inactive and the animal becoming active based on the number of times this movement was shown (up to two consecutive times: classified as still being inactive, more than two consecutive times: classified as becoming active).

**Table 2.**
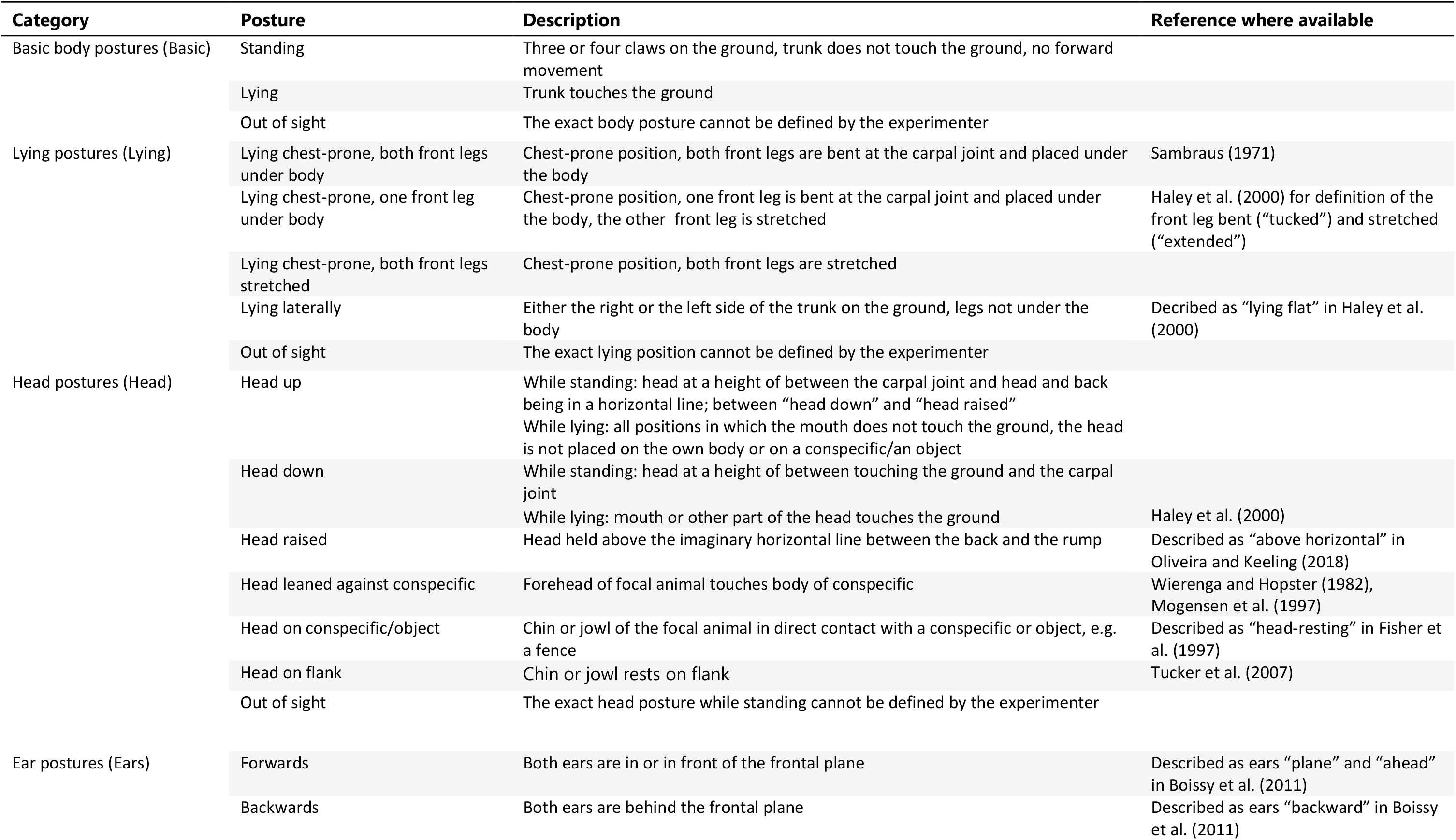
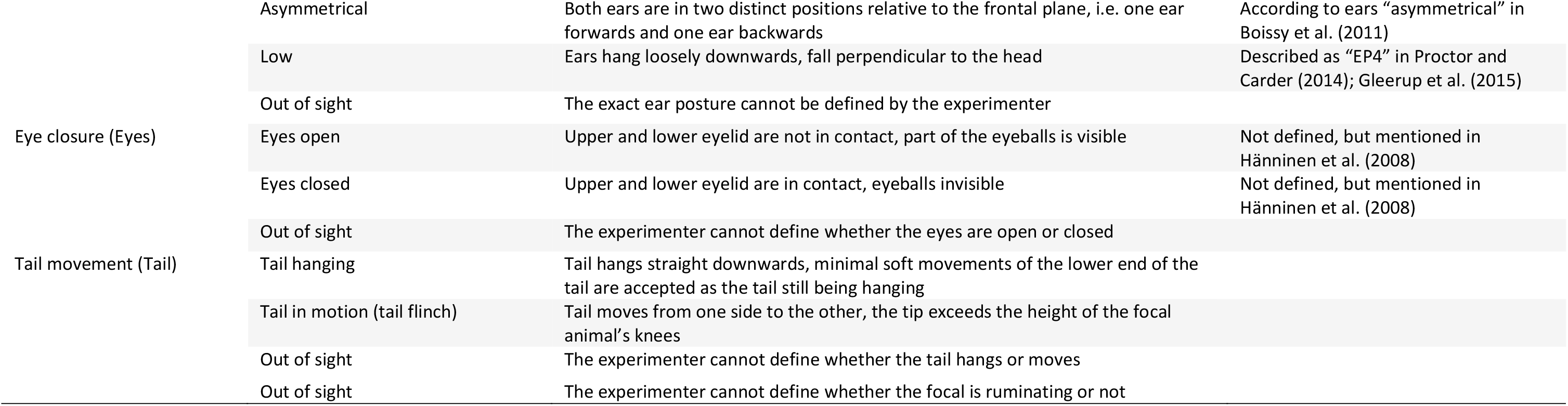
Inactivity Ethogram for fattening cattle

**Table 3.**
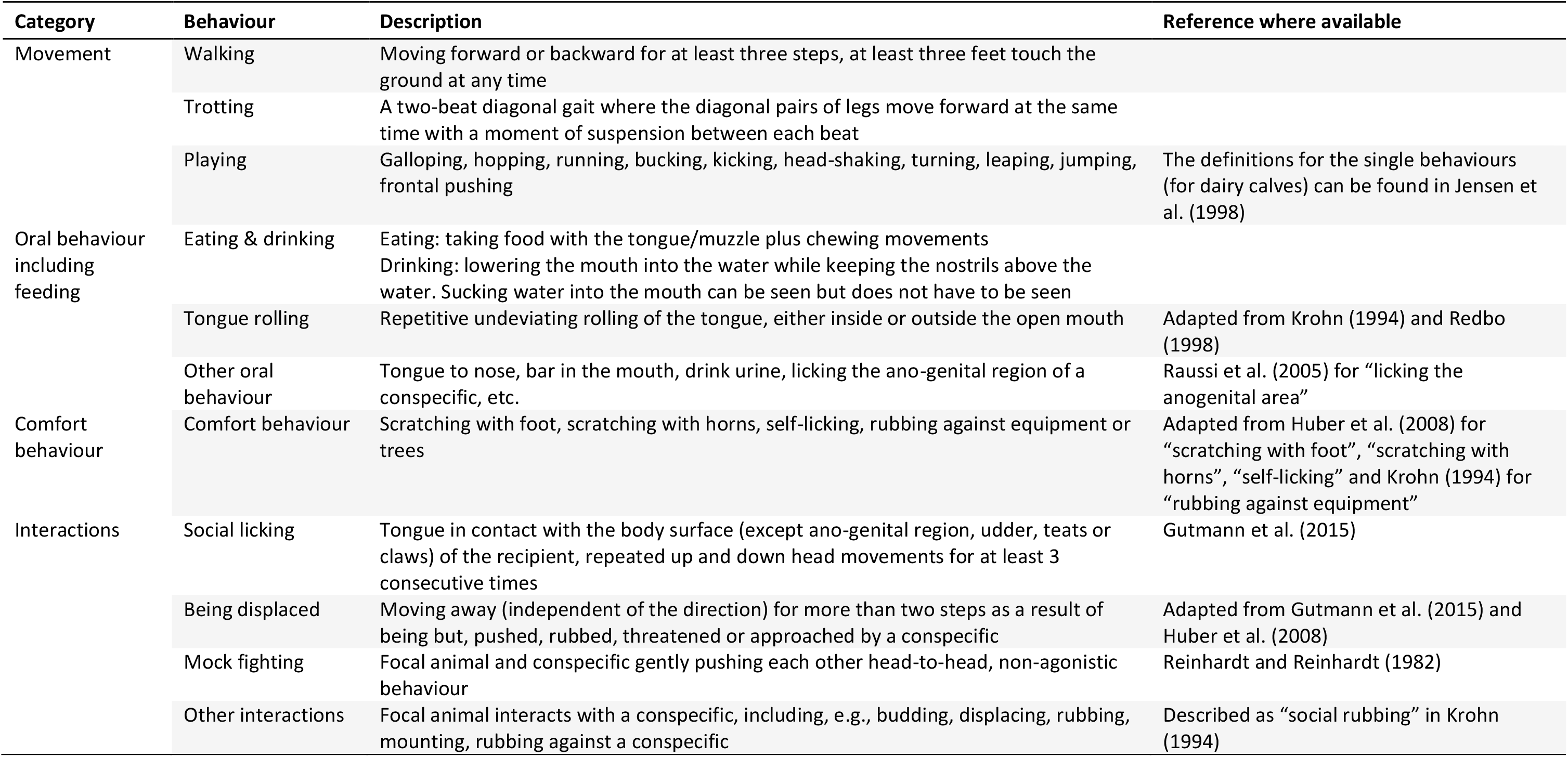
Active behaviour recorded during focal animal observations

### 2.) Application of the Inactivity Ethogram on farms with different husbandry systems

#### Experimental design

The Inactivity Ethogram was applied on nine farms in Lower and Upper Austria between April and August 2018. We visited three farms per husbandry system (intensive, semi-intensive, pasture) and all farms were visited twice, resulting in a total of 18 farm visits. Data were recorded between 9:30 AM and 06:00 PM, with one visit per farm taking place in the morning and the other one in the afternoon. The order of farm visits was counterbalanced across husbandry system and time of visit (morning/afternoon).

During each farm visit, we took 16 video clips of 15 minutes each with a Panasonic HDC-SD99 or a JVC GZ-R410BEU camcorder mounted on a tripod. Whenever possible, the 16 video clips were taken from 16 different pens on intensive and semi-intensive farms; on farms with fewer than 16 pens, we first recorded all existing pens before starting with the first pen again. The order of recorded pens was reversed during the second visit. After recording of the first eight pens, there was a break of approximately one hour before the second half of the pens was recorded. On farms with pasture, the same group of cattle was recorded for the same duration as on intensive and semi-intensive farms, i.e. for a total of four hours.

Simultaneously to the camera recording, one focal animal of the recorded pen/group was continuously observed for 15 minutes. For a focal animal to be chosen, it had to be defined as being inactive for at least 30 seconds prior to the start of observation. Moreover, focal animals were selected based on predefined rules to avoid a biased selection of animals by balancing for distance between focal animal and observer and for the position of the focal animal within its group. Data were recorded live using Mangold INTERACT^®^ (light version 17.1.11.0) on a Microsoft tablet (Acer Iconia W510). Live observations were done because they enabled us to record more of the details we were interested in, e.g. whether an animal’s eyes were open or closed, which we could not detect on the video clips taken during our pilot visits. Previous studies of inactivity did live observations as well, e.g. in mink (Meagher et al., 2017; Meagher and Mason, 2012), mice (Fureix et al., 2016), horses (Fureix et al., 2012) and in a study recording ear, neck and tail postures in dairy cattle (de Oliveira and Keeling, 2018). During the observations, the observer stood on a ladder (feet approximately 80 cm above the ground) to have a better view into the pen, e.g. for focal animals further back in the pen or on pasture. Whenever needed, especially on pasture, she used a binocular.

#### Farms

We visited three farms with fully-slatted floor pens (INTENSIVE), three farms with a feeding area, an activity area and a lying area bedded with straw (SEMI) and three farms where the animals were kept on pasture (PASTURE); one farm with daytime pasture and two farms with access to the pasture during day and night (Figure 1 and Table 4).

**Table 4.** Overview of some farm characteristics per husbandry system

|  | INTENSIVE | SEMI | PASTURE |
| --- | --- | --- | --- |
| Separate areas | no | yes: feeding area, activity area, lying area | structured by trees and location of the drinker(s) |
| Floor type | fully-slatted floor | deep-littered lying area and slatted or solid floor in feeding area | grass/soil |
| Straw bedding | no | yes | no |
| Outdoor run | no | no | n.a. |
| Number of animals per group | 4 – 10 | 4 – 14 | 9 - 31 |
| Space allowance per animal | 2.9 m<sup>2</sup><br>(2.2 – 4.2 m<sup>2</sup>) | 3.9 m<sup>2</sup><br>(2.9 – 5.5 m<sup>2</sup>) | ~ 1250 m<sup>2</sup><br>(250 – 3800 m<sup>2</sup>) |
| Feeding system | total mixed ration | total mixed ration | grass plus total mixed ration (2 farms with day grazing only); grass only on the farm with continuous pasturing |

**Figure 1.**
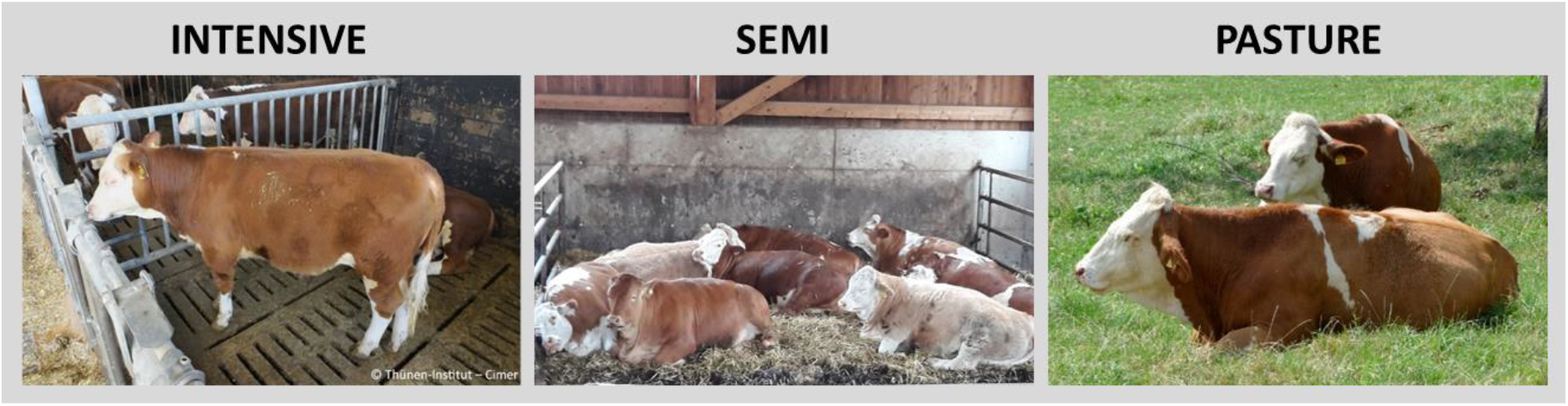
Pictures illustrating the three husbandry systems. INTENSIVE: heifers in pens with fully-slatted floor. SEMI: heifers in a straw-bedded lying area. PASTURE: heifers on pasture.

#### Animals

All focal animals were female Austrian Fleckvieh or Fleckvieh crosses, while non-focal animals were sometimes of different breeds (e.g. Limousin, Belgian Blue) and crossbreeds. We only observed heifers since bulls are very rarely kept on pasture in Austria and we aimed to avoid confounding sex and husbandry system. Heifers ranged between 8 and 27 months in age and weighed at least 300 kg according to the farmers’ estimations.

### Data preparation for analysis

#### Group level data from the video recordings

Inactivity on the pen/group level was analysed in all 288 15-minute video clips (18 farm visits × 16 video clips per farm). Scan samples were conducted on still images at minutes 2, 7 and 12, and the number of inactive animals as well as the number of inactively lying animals was recorded (n = 858 scans, 288 video clips × 3 scans per clip with six scans missing due to camera failure, “group sample datasheet”). An animal was recorded as inactive if it fulfilled the behavioural criteria for being inactive as described above for at least 30 seconds previous to the time when the still image was analysed to ensure that the same definition of inactivity was applied on both the group and the individual level. Video clips were analysed in random order by an observer who was not involved in this study otherwise. She was blind with respect to the farm, the number of visit (first or second) and the time of visit (morning or afternoon), but not to the husbandry system which was identifiable on the video clips.

#### Focal animal data from the live observations

Data from the live observations were exported from Mangold INTERACT^®^ in two different formats. For a descriptive overview of the data, the total duration of being inactive, per posture while being inactive and per active behaviour was exported for each animal (n = 288, “individual duration datasheet”). For inferential analysis, data of all postures while being inactive were extracted per second, i.e. that the continuously recorded data were translated to one-second interval samples. As a result, we received information of the postures per body part and sampling point (i.e. seconds), allowing us to analyse which postures occurred simultaneously. Only sampling points that yielded complete information of the Basic body postures, Head, Ear, Eye and Tail were considered. Sampling points for which one or more of the postures had been recorded as “out of sight” were discarded (“individual sample datasheet”).

### Statistical analyses

#### Scan samples on the group level

For a descriptive overview, group-level data are presented as the mean percentage of animals being inactive and of animals lying while being inactive per pen/group and per sampling point (n = 858 sampling points). The effect of the husbandry system on the two outcome measures was analysed with linear mixed-effects models (package: nlme, function: lme; Pinheiro et al., 2016) and generalised linear mixed-effects models (package: lme4, function: glmer, family: binomial; Bates et al., 2014) using R (version 3.6.0). Husbandry system (INTENSIVE, SEMI, PASTURE) was used as fixed effect, and random effects were pen (1 - 16, three sampling points per pen/group) nested in farm visit (1 - 2) nested in farm (1 - 9). Animals kept on pasture were all from one group per farm, which is why we did not have a pen level for PASTURE. However, since observations were always interrupted for approximately one hour after the first eight focal animals had been observed, we differentiated between the first block of observations (animals 1 – 8) and the second block of observations (animals 9 – 16) to account for variation at the group level for the three PASTURE farms. Model assumptions for the linear mixed-effects models (normal distribution, homogeneity of variance) were verified by graphical inspection of the residuals and data were converted to a binary measure (0,1) when necessary. This was the case for “lying” for which we converted the proportion of lying animals per sampling point to 0/1 with 0 meaning that no animal was lying and 1 meaning that at least one animal was lying. Data were then analysed with generalised linear-mixed effects models (see Table 5 for details per outcome measure). Statistical significance was considered to be p < 0.05 for all (generalised) linear mixed-effects models.

**Table 5.**
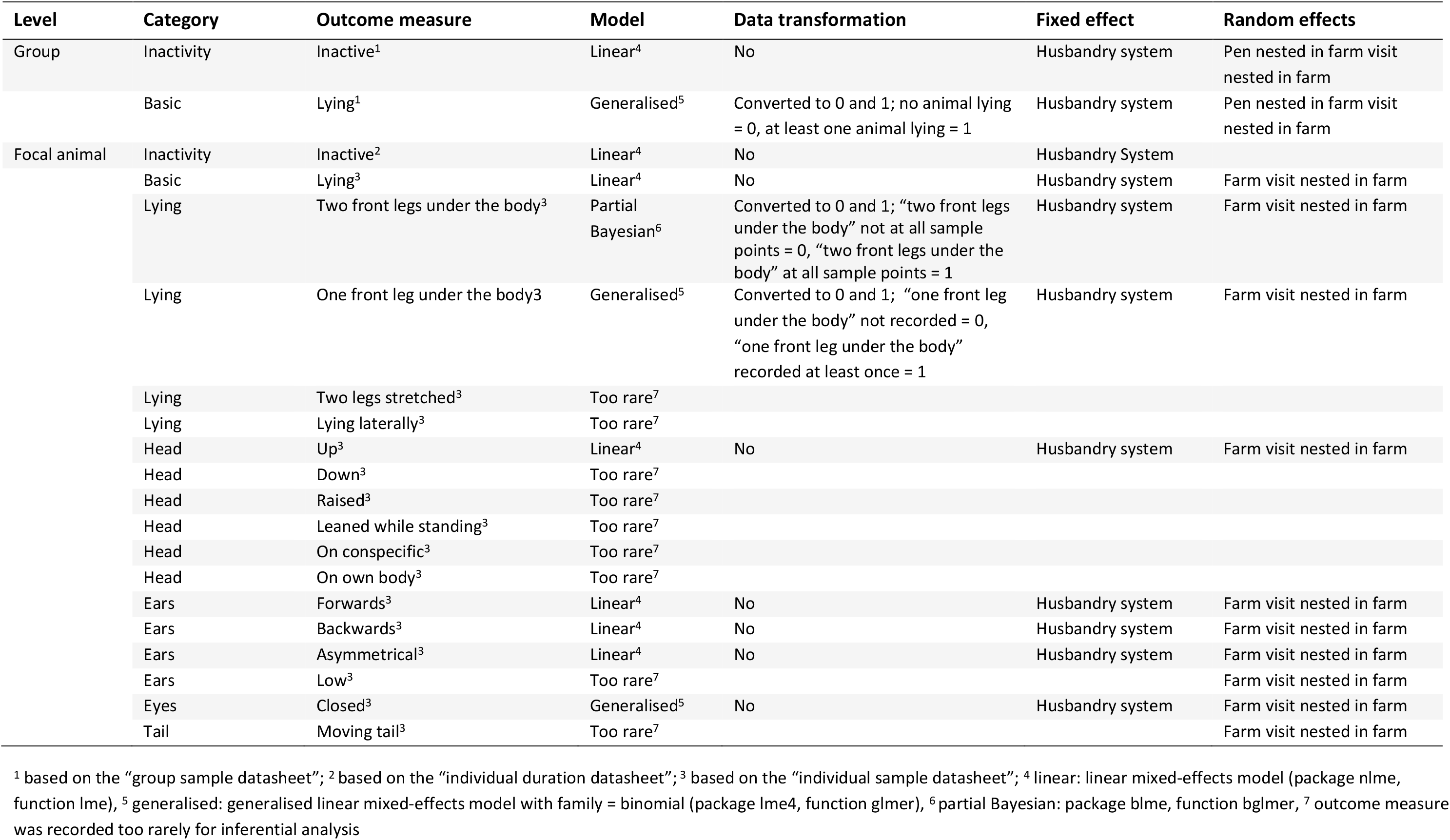
Overview of the statistical models, fixed and random effects for the outcome measures on the percentage of time of the single postures

| Level | Category | Outcome measure | Model | Data transformation | Fixed effect | Random effects |
| --- | --- | --- | --- | --- | --- | --- |
| Group | Inactivity | Inactive<sup>1</sup> | Linear<sup>4</sup> | No | Husbandry system | Pen nested in farm visit nested in farm |
|  | Basic | Lying<sup>1</sup> | Generalised<sup>5</sup> | Converted to 0 and 1; no animal lying = 0, at least one animal lying = 1 | Husbandry system | Pen nested in farm visit nested in farm |
| Focal animal | Inactivity | Inactive<sup>2</sup> | Linear<sup>4</sup> | No | Husbandry System |  |
|  | Basic | Lying<sup>3</sup> | Linear<sup>4</sup> | No | Husbandry system | Farm visit nested in farm |
|  | Lying | Two front legs under the body<sup>3</sup> | Partial Bayesian<sup>6</sup> | Converted to 0 and 1; “two front legs under the body” not at all sample points = 0, “two front legs under the body” at all sample points = 1 | Husbandry system | Farm visit nested in farm |
|  | Lying | One front leg under the body<sup>3</sup> | Generalised<sup>5</sup> | Converted to 0 and 1; “one front leg under the body” not recorded = 0, “one front leg under the body” recorded at least once = 1 | Husbandry system | Farm visit nested in farm |
|  | Lying | Two legs stretched<sup>3</sup> | Too rare<sup>7</sup> |  |  |  |
|  | Lying | Lying laterally<sup>3</sup> | Too rare<sup>7</sup> |  |  |  |
|  | Head | Up<sup>3</sup> | Linear<sup>4</sup> | No | Husbandry system | Farm visit nested in farm |
|  | Head | Down<sup>3</sup> | Too rare<sup>7</sup> |  |  |  |
|  | Head | Raised<sup>3</sup> | Too rare<sup>7</sup> |  |  |  |
|  | Head | Leaned while standing<sup>3</sup> | Too rare<sup>7</sup> |  |  |  |
|  | Head | On conspecific<sup>3</sup> | Too rare<sup>7</sup> |  |  |  |
|  | Head | On own body<sup>3</sup> | Too rare<sup>7</sup> |  |  |  |
|  | Ears | Forwards<sup>3</sup> | Linear<sup>4</sup> | No | Husbandry system | Farm visit nested in farm |
|  | Ears | Backwards<sup>3</sup> | Linear<sup>4</sup> | No | Husbandry system | Farm visit nested in farm |
|  | Ears | Asymmetrical<sup>3</sup> | Linear<sup>4</sup> | No | Husbandry system | Farm visit nested in farm |
|  | Ears | Low<sup>3</sup> | Too rare<sup>7</sup> |  |  | Farm visit nested in farm |
|  | Eyes | Closed<sup>3</sup> | Generalised<sup>5</sup> | No | Husbandry system | Farm visit nested in farm |
|  | Tail | Moving tail<sup>3</sup> | Too rare<sup>7</sup> |  |  | Farm visit nested in farm |

<sup>1</sup> based on the “group sample datasheet”; <sup>2</sup> based on the “individual duration datasheet”; <sup>3</sup> based on the “individual sample datasheet”; <sup>4</sup> linear: linear mixed-effects model (package nlme, function lme), <sup>5</sup> generalised: generalised linear mixed-effects model with family = binomial (package lme4, function glmer), <sup>6</sup> partial Bayesian: package blme, function bglmer, <sup>7</sup> outcome measure was recorded too rarely for inferential analysis

#### Continuous observations on the focal animal level

##### a) Percentage of time the single postures were observed for

The analysis of the total time the animals spent inactive was based on the “individual duration datasheet”, while all other outcome measures were analysed on the basis of the “individual sample datasheet”. We divided the number of sampling points (i.e. seconds) for each posture (e.g. “Ears forward”) by the total number of sampling points for the respective body part (i.e. all sampling points for which Ears were recorded). This allowed to account for differences between animals in the number of sampling points (since the postures were only recorded when the focal animal was inactive) and for the number of sampling points the respective body part (e.g. Ears) was recorded as “out of sight”. The effect of husbandry system on the different postures shown by the focal animals was analysed in the same way as for the group-level data, but random effects on the focal animal level were farm visit (1 – 2) nested in farm (1 - 9) without considering the pen, since we did not have repeated measures and thus no variation on this level. For the outcome measure “Two front legs under the body”, we used a partial Bayesian method (package: blme, function: bglmer; Dorie, 2011) due to a singular fit using a glmer model. The models for each outcome measure, conversion to 0/1 when necessary, and fixed as well as random effects are given in Table 5. As a measure of between-farm variation, we calculated the Variance Partitions Coefficients (VPC) of the random effect farm by dividing the variance coefficient of farm by the total variance of all random effects (set to 1 for glmer and blmer models) for the analysed outcome measures (Goldstein et al., 2010).

##### b) Simultaneous occurrences of postures of different body parts

The analyses of the simultaneous occurrences of postures were based on the “individual sample datasheet”. We included whether the animals were standing or lying (Basic), their head postures (Head), ear postures (Ears), and the closure of the eyes (Eyes). Tail movement was excluded since tail was recorded as “hanging” for most of the time and was in motion almost exclusively on PASTURE. To understand patterns in simultaneous observation of postures in a manner which accounts for frequency of occurrence within as well as between animals, a machine learning algorithm was used. The **S**equential **PA**ttern **D**iscovery using **E**quivalence classes (spade) algorithm (Zaki, 2001) was used for this purpose in R (package: arulesSequences, function: cspade; Buchta and Hahsler, 2019). This algorithm is commonly used to identify which items are purchased together in shopping baskets or in subsequent shopping trips (Koenecke, 2019). When using this algorithm, the support threshold was set at 0.1 and the maximum gap (maxgap) was set at 0 to search only for postures that occurred simultaneously (and not in subsequent observations). Support is the proportion of a given combination out of the total number of co-occurring combinations. The support threshold is the value above which co-occurring combinations are identified as frequent combinations. The algorithm outputs the confidence for given co-occurring combinations, which is the likelihood of observing that combinations, in this case the combination of body parts, are displayed in future observations. This is calculated by the conditional probability of observing the combination given it has already been observed in that animal. The cspade algorithm was calculated separately for each husbandry system. To account for the fact that the different husbandry systems had different number of total observations and different number of animals, we compared cspade results from full data to data truncated to account for this (reduced so that husbandry systems matched for number of observations and number of animals in a randomised manner). Since truncated data and full datasets were not different (correlation r > 0.99), we present data from the full datasets here. The confidence of pairwise combinations of body parts (frequent sequences of two) and all four body parts is displayed, the former as a network figure with confidence used as the strength of connections between body parts (using visNetwork and igraph). Differences in confidence noted between husbandry systems are presented and since several studies defined animals as being inactive if they were immobile with eyes open (Meagher and Mason, 2012; Fureix et al. 2016), we also focus on presenting the confidence values for the pairwise combinations “Lying with Eyes open”, “Lying with Eyes closed”, “Standing with Eyes open” and “Standing with Eyes closed”.

## Results

### Scan samples on the group level

The percentage of inactive animals per pen/group was lowest on PASTURE (mean ± standard deviation: 14.7 % ± 0.24), intermediate on SEMI (25.0 % ± 0.25) and highest on INTENSIVE farms (35.4 % ± 0.28), but did not differ statistically significant between husbandry systems (F_2,6_ = 2.57, p = 0.16; Figure 2 and Table 6). When being inactive, the percentage of lying animals per group was highest on PASTURE (74.4 % ± 0.41), followed by SEMI (65.94 % ± 0.40) and INTENSIVE farms (63.8 % ± 0.43), but again there was no statically significant difference between husbandry systems (X^2^_2_ = 0.63, p = 0.73).

**Table 6.**
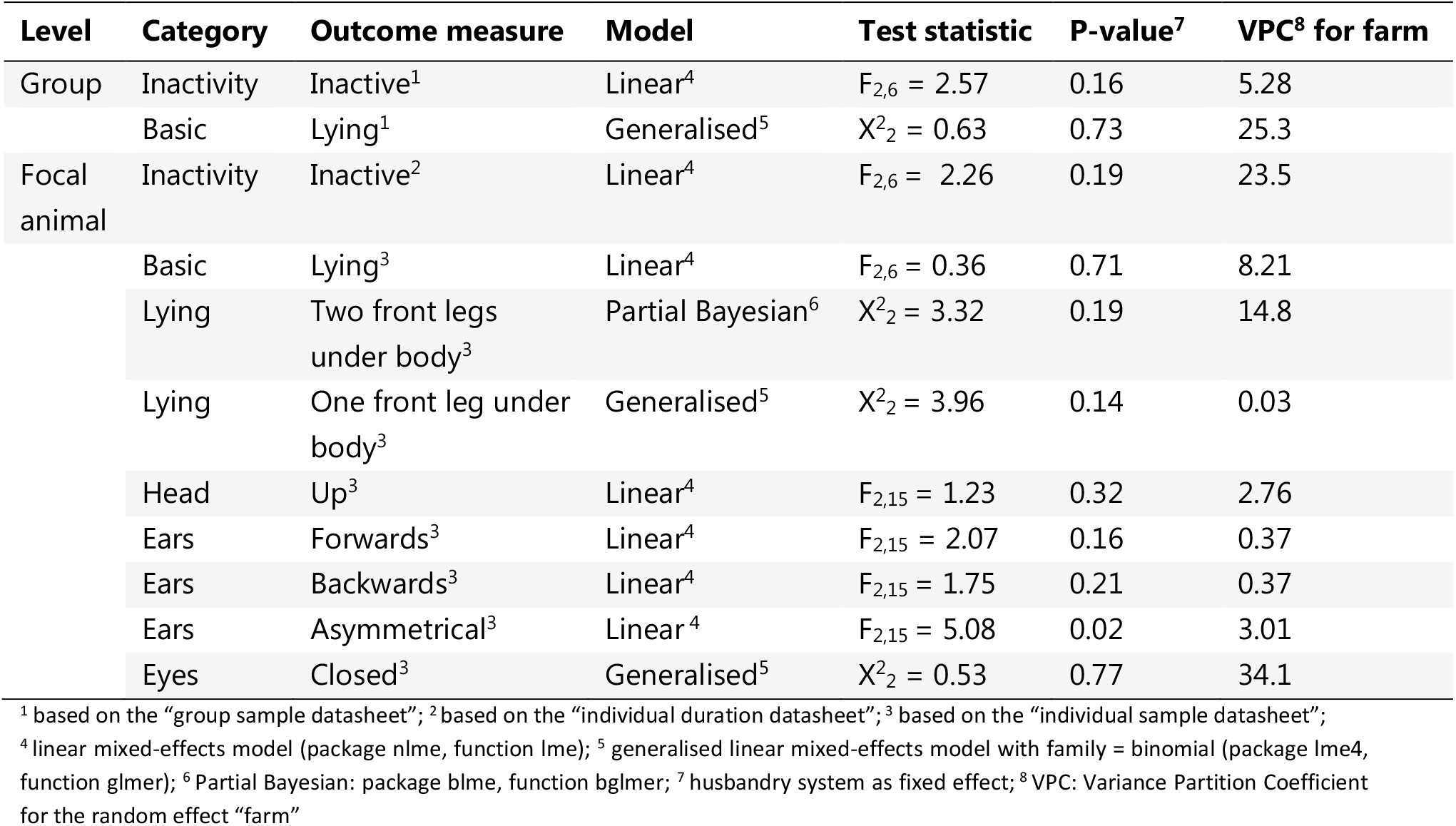
Results of the outcome measures that were analysed statically

**Figure 2.**
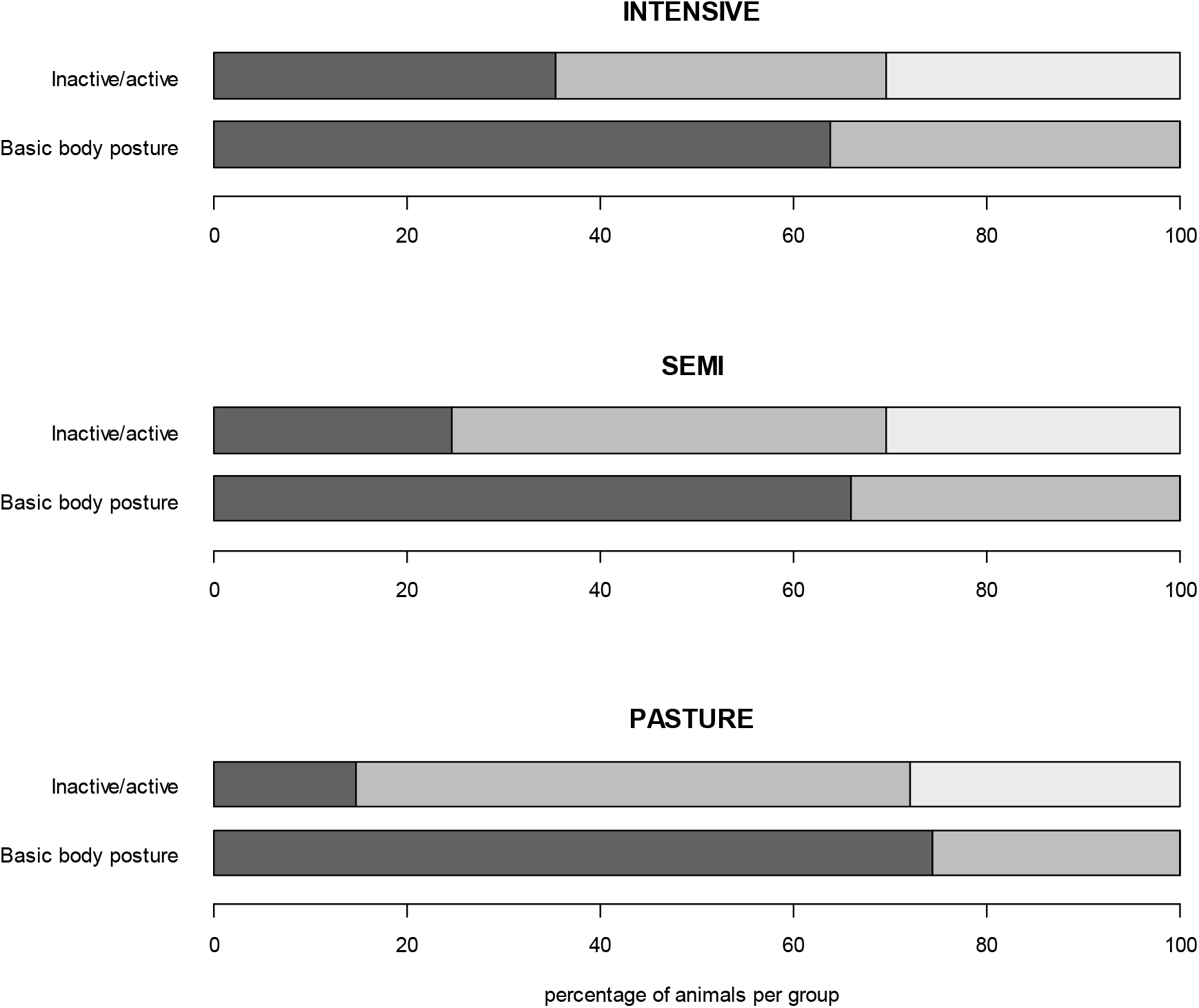
(In)activity levels and basic body postures on the group level. The mean percentage of inactive animals per group and of inactive animals lying or standing is shown for the three different husbandry systems. **Inactive/active**: inactive (dark), active (medium), out of sight (light). **Basic body posture**: lying (dark), standing (medium) while being inactive.

### Continuous observations on the focal animal level

#### Percentage of time the single postures were observed for

The pattern of the focal animals’ inactivity level corresponded to the inactivity level on the group level with cattle spending least time inactively on PASTURE (55.8 % ± 40.0), followed by SEMI (75.0 % ± 27.1) and INTENSIVE farms (85.5 % ± 20.7), but again the inactivity level did not differ significantly between husbandry systems (F_2,6_ = 2.26, p = 0.19; Figure 3 and Table 6). Compared to the results from the group level data, inactive focal animals spent less time lying (50.8 % ± 47.1) and thus more time standing, and while the time the animals spent lying followed the increasing intensity of the husbandry system on the group level, it slightly decreased from INTENSIVE (57.4 % ± 47.5) to SEMI (52.6 % ± 47.2) to PASTURE farms (42.1 % ± 45.7) on the focal animal level. When lying, a chest-prone position with the two front legs tucked under the body was observed for the longest (86.9 % ± 26.8 across husbandry systems) and the most commonly observed head position across all husbandry systems was the head held up (84.5 % ± 20.3). Cattle on INTENSIVE and SEMI farms showed ears backwards for the longest (INTENSIVE: 48.2 % ± 27.3; SEMI: 44.0 % ± 28.3), followed by ears forward (INTENSIVE: 32.8 % ± 27.5; SEMI: 37.3 % ± 29.4), while these two ear postures were displayed almost equally long by cattle on PASTURE (backwards: 38.1 % ± 30.8, forwards: 38.6 % ± 31.3) The percentage of time cattle displayed low ears increased from INTENSIVE (2.4 % ± 8.4) to SEMI (4.2 % ± 12.2) to PASTURE (15.2 % ± 27.3), whereas the time their ears were in an asymmetrical position increased with increasing intensity of the system (PASTURE: 3.9 % ± 12.3, SEMI: 9.5 % ± 12.0, INTENSIVE: 12.4 % ± 14.1; p = 0.02 as based on the “individual sample datasheet”). On INTENSIVE and SEMI farms, the animals’ eyes were open for approximately four fifths of the observed time (INTENSIVE: 81.7 % ± 26.6, SEMI: 81.4 % ± 30.6), while they were open for approximately 70 % of the inactivity time in cattle on PASTURE (69.9 % ± 38.4). As described above, the tail was hanging for almost all of the inactivity time (more than 99 % on INTENSIVE and SEMI farms), while it was recorded as moving for 13.1 % (± 24.7) of the time in cattle on PASTURE. None of the outcome measures was statistically significant influenced by the husbandry system (all p-values > 0.05), except for the outcome measure “Ears asymmetrical” (Table 6).

**Figure 3.**
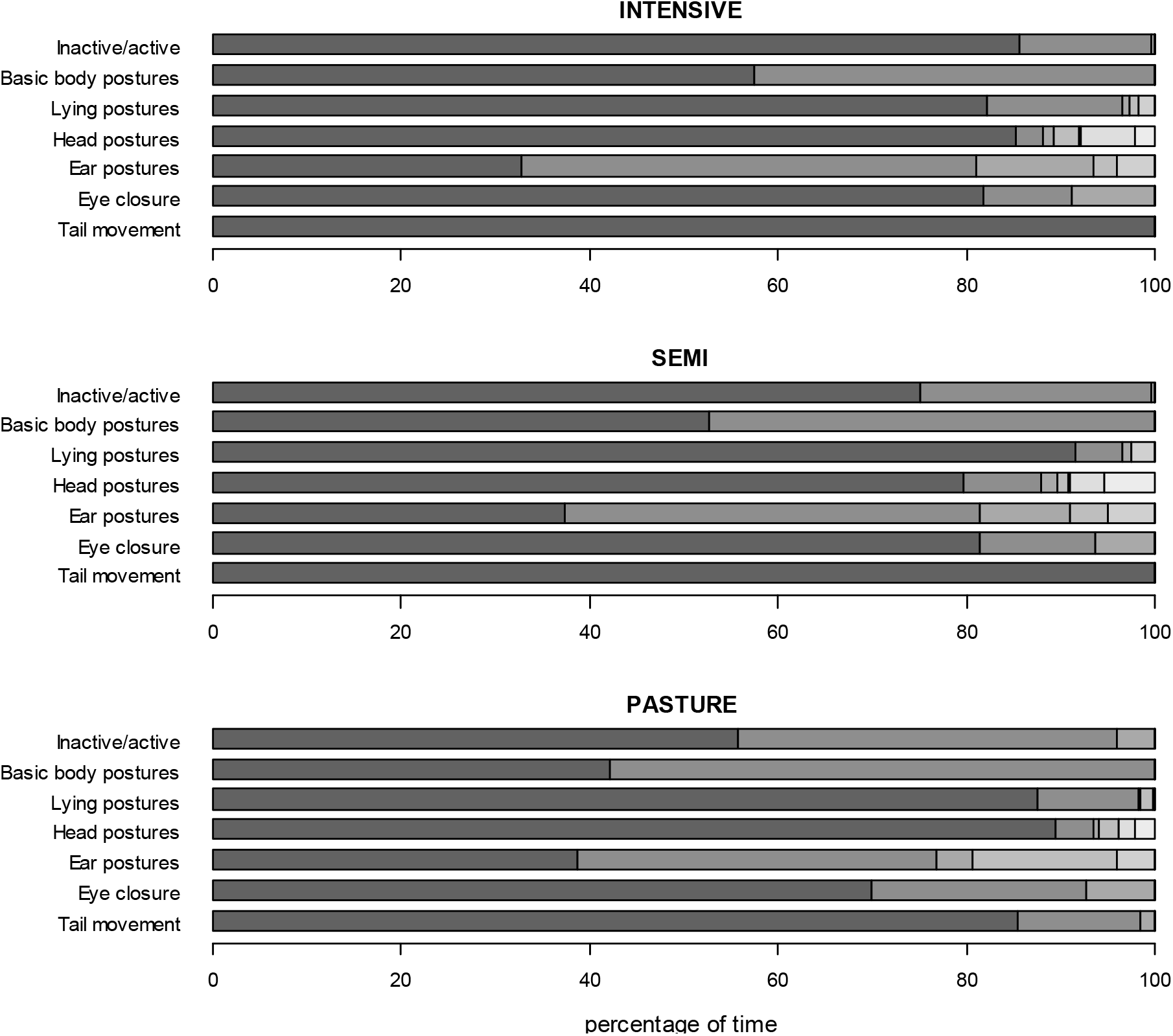
Mean percentage of time the focal animals showed the different postures while being inactive. Postures per category from left to right; **Inactive/active:** inactive, active, out of sight. **Basic body postures**: lying, standing. **Lying postures**: two front legs under body, one front leg under body, both front legs stretched, lateral, out of sight. **Head postures**: up, down, raised, on conspecific, leaned, on own body while lying, out of sight. **Ear postures**: forwards, backwards, asymmetrical, low, out of sight. **Eye closure**: open, close, out of sight. **Tail movement**: no movement, movement, out of sight.

An overview of the mean percentage of time animals spent performing different active behaviours per husbandry systems is given in Figure 4.

**Figure 1.**
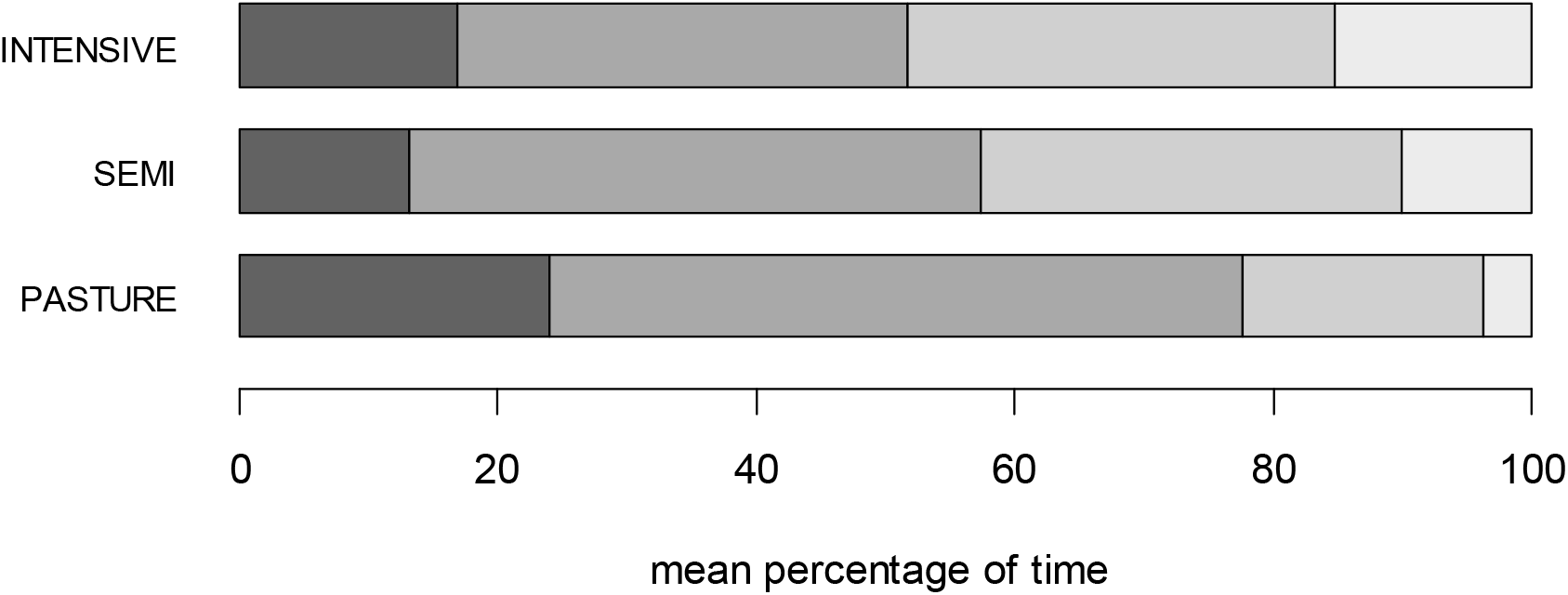
Mean percentage of time the focal animals displayed different active behaviours. For all three husbandry systems from left to right: **movement** (dark), **oral behaviour** including feeding (dark medium), **comfort behaviour** (light medium), **interactions** with conspecifics (light).

#### Variation Partition Coefficient (VPC)

Between-farm variation varied greatly between outcome measures ranging from a low VPC of 0.03 for “Lying with one front leg under the body” to a rather high VPC of 34.1 for “Eyes closed” (Table 6).

#### Simultaneous occurrence of postures of different body parts

##### a) Overview

For analyses of the simultaneous occurrences of different postures, the final dataset comprised of 131,791 time points (i.e. seconds), corresponding to 36.6 hours of observation, for which we had information of all four body parts (Basic, Head, Ears, Eyes; tail movements were excluded as described above). It did not include 17 animals for which full information of all four body parts was not available (INTENSIVE: n = 0, SEMI: n = 1, PASTURE: n = 16), leading to a total of 271 animals in the final dataset. The number of data points per animal varied greatly between individuals and was highest in INTENSIVE, followed by SEMI and PASTURE (INTENSIVE: 565 ± 268, SEMI: 451 ± 291, PASTURE: 434 ± 337).

To ensure that data were not skewed by single individuals displaying the same combination of postures for a long time, we used the cspade algorithm which accounts for the frequency of occurrences both within and between animals and used the confidence, i.e. the likelihood of observing the respective combination of postures displayed in future observations, from this algorithm as outcome measure (ranging from 0 to 1). Between 75 (PASTURE) and 92 (SEMI, INTENSIVE) posture combinations were identified as co-occurring frequently using the cspade algorithm (Table 7).

**Table 7.** Number of frequently co-occurring body parts as identified by the cspade algorithm

|  | **Two body parts** | **Three body parts** | **Four body parts** | **Total of combinations** |
| --- | --- | --- | --- | --- |
| INTENSIVE | 39 | 40 | 13 | 92 |
| SEMI | 37 | 40 | 15 | 92 |
| PASTURE | 32 | 32 | 11 | 75 |

##### b) Pairwise combinations of postures

The most frequently co-occurring combinations of postures of two body parts (e.g. Basic plus Ear, Head plus Eye) across individuals were similar between husbandry systems (Figure 5). The combination with the highest confidence was “Head up with Eyes open” for all husbandry systems. A small number of combinations were identified as frequently occurring in one or two husbandry system only. For example, “Ears low” co-occurred with “Lying”/“Head up”/“Eyes open” and “Eyes closed” frequently enough to be displayed only on PASTURE, co-occurred with the same postures but not with “Eyes closed” in SEMI and did not co-occur with any posture in INTENSIVE systems. Conversely, “Head raised” only co-occurred with other postures in INTENSIVE and SEMI systems, but not on PASTURE. These combinations tended to be those with the lowest confidence values, which were just above the cut-off threshold. “Lying with Eyes open” as a combination had a lower confidence on PASTURE, particularly when compared to INTENSIVE, whereas it was the opposite for “Standing with Eyes open” (Table 8).

**Table 8.** Confidence values for Lying and Standing with Eyes open and Eyes closed

|  | Lying with<br>Eyes open | Lying with<br>Eyes closed | Standing with<br>Eyes open | Standing with<br>Eyes closed |
| --- | --- | --- | --- | --- |
| INTENSIVE | 0.61 | 0.40 | 0.52 | <0.01 |
| SEMI | 0.59 | 0.32 | 0.53 | 0.05 |
| PASTURE | 0.55 | 0.38 | 0.64 | <0.01 |

Confidence values (likelihood of observing the respective combination of postures displayed in future observations) can range between 0 (not likely) and 1 (very likely).

**Figure 5.**
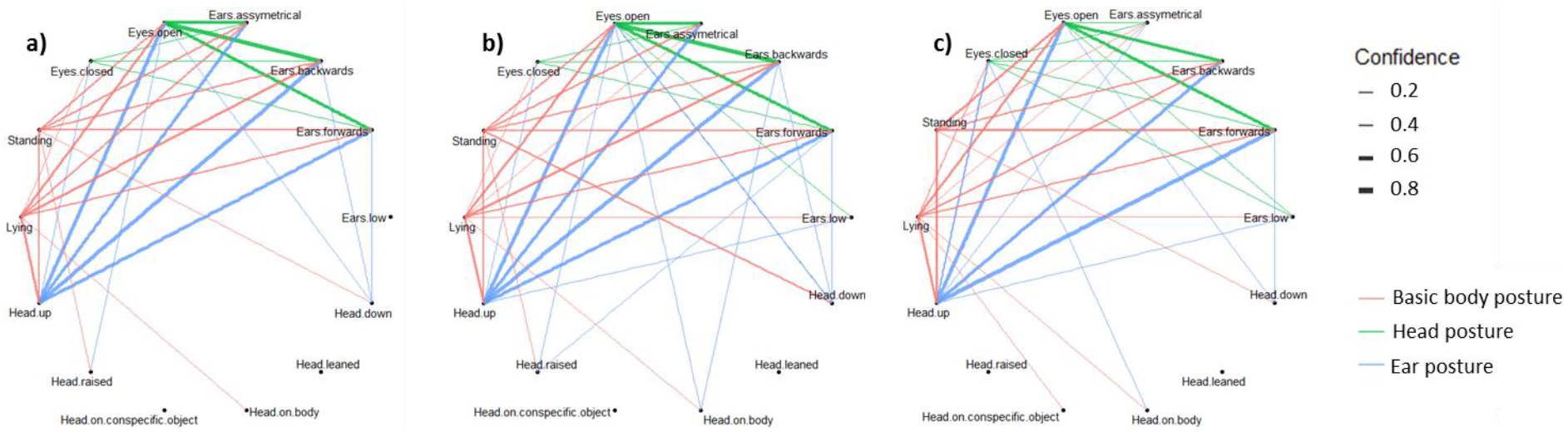
Confidence of pairwise combinations of postures. The confidence of pairwise combinations of postures (frequent sequences of two) of Basic, Head, Ears and Eyes displayed for a) INTENSIVE, b) SEMI and c) PASTURE husbandry systems. The thickness of the lines indicates the confidence values of the combination of postures calculated using the cspade algorithm. The colours indicate the involved body parts; red: Basic plus Head, Ear or Eye; green: Head plus Ear or Eye; blue: Ear plus Eye.

##### c) Fourfold combinations of postures

The fourfold combination shown for the largest percentage of time in INTENSIVE and SEMI systems was “Lying with Head up, Ears backwards and Eyes open”, while it was “Lying with Head up, Ears low and Eyes closed” on PASTURE (Figure 6). Of the frequently occurring fourfold combinations of Basic, Head, Ears and Eyes, ten were the same across all husbandry systems (confidence ≥ 0.1 for all three husbandry systems, Table 9).

**Table 9.** Confidence values by husbandry system for frequently occurring combinations of postures of all four body parts (Basic, Head, Ears, Eyes) identified using the cspade algorithm

| Body part combination | INTENSIVE | SEMI | PASTURE |
| --- | --- | --- | --- |
| Lying_Head up_Ears backwards_Eyes open | 0.604 | 0.568 | 0.363 |
| Standing_Head up_Ears forwards_Eyes open | 0.490 | 0.484 | 0.575 |
| Standing_Head up_Ears backwards_Eyes open | 0.479 | 0.400 | 0.425 |
| Lying_Head up_Ears forwards_Eyes open | 0.438 | 0.442 | 0.375 |
| Standing_Head up_Ears asymmetrical_Eyes open | 0.417 | 0.411 | 0.175 |
| Lying_Head up_Ears asymmetrical_Eyes open | 0.406 | 0.326 | 0.088 |
| Lying_Head up_Ears backwards_Eyes closed | 0.333 | 0.284 | 0.338 |
| Standing_Head down_Ears forwards_Eyes open | 0.198 | 0.295 | 0.200 |
| Lying_Head up_Ears forwards_Eyes closed | 0.188 | 0.200 | 0.275 |
| Standing_Head down_Ears backwards_Eyes open | 0.188 | 0.211 |  |
| Lying_Head up_Low ears_Eyes closed |  | 0.105 | 0.200 |
| Lying_Head up_Low ears_Eyes open |  | 0.116 | 0.163 |
| Lying_Head up_Ears asymmetrical_Eyes closed | 0.156 | 0.126 | 0.138 |
| Lying_Head on body_Ears backwards_Eyes open |  | 0.137 |  |
| Standing_Head raised_Ears forwards_Eyes open | 0.115 | 0.126 |  |
| Lying_Head on conspecific or object_Ears backwards_Eyes open | 0.104 |  |  |

Only combinations with a confidence value $\geq 0.1$ are shown. Combinations are ordered from the largest to the smallest maximum confidence value across husbandry systems. For each combination, the largest confidence value is marked in dark blue, the second smallest in medium blue, the smallest in light blue and values less than 0.1 in white.

**Figure 6.**
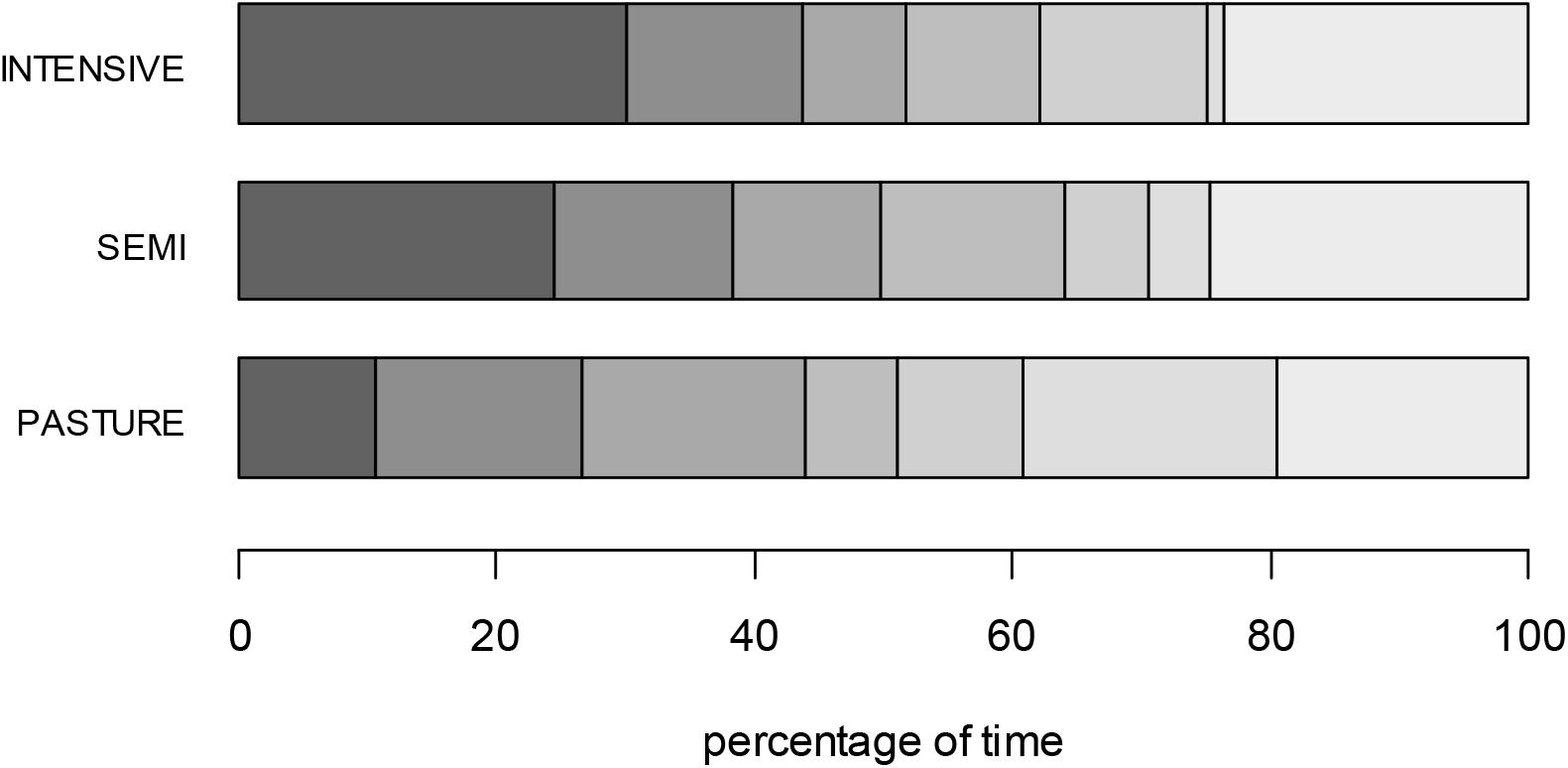
Mean percentage of time the animals spent in the six most common fourfold combinations of postures (combinations displayed for at least 5 % of the inactivity time across husbandry systems). For all three husbandry systems from left to right: **(1)** Lying_Head up_Ears backwards_Eyes open, **(2)** Standing_Head up_Ears forwards_Eyes open, **(3)** Lying_Head up_Ears backwards_Eyes closed, **(4)** Lying_Head up_Ears forwards_Eyes open **(5)** Standing_Head up_Ears backwards_Eyes open, **(6)** Lying_Head up_Ears low_Eyes closed, **(7)** Other combinations

## Discussion

The aims of our study were to develop an Inactivity Ethogram for fattening cattle and to apply this ethogram on farms with different husbandry systems to explore what inactive behaviour in cattle looks like. Scan samples of the groups and continuous observations of the focal animals revealed no statistically significant differences between husbandry systems with respect to the animals’ overall inactivity level and the time the different postures were observed for while the animals were inactive (except for Ears asymmetrical, which were more prevalent in INTENSIVE than in SEMI than in PASTURE). Using the cspade algorithm, we detected frequently co-occurring postures of different body parts, thereby identifying differences in the patterns of postural combinations across husbandry systems and providing further insight into the complexity of inactive behaviour in fattening cattle.

Our Inactivity Ethogram encompasses various postures of different body parts and may be a promising methodological tool in future studies investigating different kinds of inactive behaviours and what they mean for animal welfare. Especially in conditions where the valence of the animals’ inactivity is specifically manipulated and/or investigated, the application of our Inactivity Ethogram may be used to detect indicators that help differentiating between positive inactivity, for example reflecting relaxation, and negative inactivity, for example reflecting boredom or depression-like states. Detailed observation of the postures of different body parts is especially useful in the study of inactive behaviour, where overt behaviour is rarely shown and subtle differences in postures that are not captured with traditional ethograms may reveal important information. However, despite its particular relevance for the study of inactive behaviour, the principle of recording details of the animals’ postures and the simultaneous occurrence of postures of different body parts may advance the study of emotional states more generally. De Oliveira and Keeling (2018), for example, already stressed the importance of capturing details of the animals’ postures, when they investigated ear and neck positions as well as tail movements of dairy cows exposed to three routine situations to identify indicators of emotional valence.

Since our Inactivity Ethogram was developed and applied on farms with different husbandry systems, it may be used in a wide range of conditions. However, we only observed Fleckvieh heifers in our study, which is why the ethogram may need to be adapted to other cattle breeds and bulls. The general principle of the ethogram structure may be transferred to other species than cattle and may be further extended. Depending on the specific research questions, rumination and vocalisations (e.g. humming, mooing) could, for example, be included in future studies.

The overall level of inactivity and the percentage of time for which each of the postures was displayed did not differ statistically significant between the three husbandry systems (with the exception for Ears asymmetrical), but the number of animals being inactive per group and the time focal animals were inactive rose with increasing intensity of the system from PASTURE to SEMI to INTENSIVE. This finding is in line with other studies that show increased inactivity levels in barren compared to enriched housing conditions in pigs (Bolhuis et al., 2006, 2005), minks (Meagher and Mason, 2012) and mice (Fureix et al., 2016). However, our study design differed from these existing studies in two main aspects. First, we did not specifically compare barren and enriched conditions, but whole housing systems that differed in many more aspects than just barrenness. Second, and related to the first aspect, housing conditions in our study differed between farms whereas the previous studies were conducted in an experimental setting, in which the conditions were manipulated within farm. Between-farm noise might thus have masked potential differences in our study, which is why we calculated the Variation Partition Coefficient (VPC) as a measure of between-farm variation. The VPC varied a lot between the different outcome measures with relatively small values for the ear postures and rather large values for the basic body postures on the group level, the overall inactive behaviour displayed by the focal animals and the closure of the eyes. However, since not all VPCs were large, between-farm variation cannot solely explain why we did not find differences between husbandry systems; of course it is also possible that husbandry system *per se* did simply not have an influence on the time the single postures were shown for.

Even though the percentage of time the single postures were observed for did not differ statistically significant between husbandry systems, our study gives insight into potentially interesting patterns. Ears held asymmetrically, for example, increased from PASTURE to SEMI to INTENSIVE (in this case statistically significant), while it was the exact opposite for the occurrence of low ears (PASTURE > SEMI > INTENSIVE). Other studies on ear postures might have served for a tentative interpretation of our findings, but, unfortunately, the results from the existing studies are quite inconsistent. For example, one of the most commonly shown ear postures in this and other studies, ears backwards, has been described as reflecting both a positive state (e.g. de Oliveira and Keeling, 2018; Proctor and Carder, 2014) and a negative state (e.g. Boissy et al., 2011 in sheep; Gleerup et al., 2015). Moreover, low ears have been described to be indicative of positive low arousal states (Proctor and Carder, 2014; here described as “ears loosely hung down”), but also as a sign of pain in dairy cattle (Gleerup et al., 2015). Since low ears in our study were mostly shown on PASTURE and the percentage of time for which they were shown decreased from PASTURE to SEMI to INTENSIVE, they were more likely indicative of a positive low arousal rather than a painful state. However, we are cautious to draw firm conclusions since we did not have *a priori* hypotheses of how husbandry system would affect the emotional valence of the animals and would thus like to avoid circular reasoning. For the asymmetrical ears, de Oliveira and Keeling (2018) differentiated between right and left asymmetry depending on which ear was pointing backward, and predicted based on their results that right ear backwards might reflect a more positively valenced state than left ear backwards, in accordance with the concept of emotional lateralisation (as reviewed in Leliveld et al., 2013). We did not differentiate between left and right ear when recording asymmetry but given that the asymmetrical ears were the only outcome measure which differed statistically significant between husbandry systems, such a differentiation may be a valuable addition to our Inactivity Ethogram in future studies.

The animals’ eyes were open for approximately 80 % of the inactivity time on INTENSIVE and SEMI farms, while they were only open for approximately 70 % of the inactivity time in cattle on PASTURE. The longer time animals on PASTURE spent with closed eyes may indicate that these animals slept more during our observations, which would be in line with findings that cattle on pasture have several episodes of rest and sleep during the day (Sambraus, 1971), while cattle kept indoors mostly sleep during the night (Ternman et al., 2019. However, it needs to be stressed that the closure of the eyes alone is not sufficient to define the animals as being asleep; several characteristics have to be fulfilled to properly differentiate between sleep and resting (Nicolau et al., 2000), but those were not recorded in our study.

Since other studies defined inactivity as being still but awake with “awake” being characterised by open eyes, we specifically investigated the pairwise combinations of Lying and Standing with either Eyes open or closed. The confidence values for “Lying with Eyes open” were largest for INTENSIVE, followed by SEMI and PASTURE systems, in line with other studies who found animals to be more still with eyes open in barren compared to enriched conditions (e.g. Fureix et al., 2016; Meagher et al., 2017; Meagher and Mason, 2012). However, the exact opposite pattern was observed for “Standing with Eyes open”, with the largest confidence value for PASTURE, followed by SEMI and INTENSIVE. This result indicates that future studies should differentiate between different basic body postures, depending on the species of interest, and the simultaneous closure of the eye. Cattle on INTENSIVE farms did not only have the largest confidence values for “Lying with Eyes open”, but also for “Lying with Eyes closed”, which is in accordance with the findings from Bolhuis and colleagues (2005) in fattening pigs that spent more time lying with both open and closed eyes in barren compared to enriched conditions. This result indicates that lying with eyes open goes along with more lying overall, but our finding needs to be interpreted carefully. First, the comparably higher value for animals from INTENSIVE farms do not necessarily mean that the same individuals accounted for both “Lying with Eyes open” and “Lying with Eyes closed”. Second, species-specific differences need to be taken into account. While being still with eyes open may be a good indicator of boredom or depression-like states in mink and mice, respectively, cattle spend about seven hours a day ruminating (Beauchemin, 2018) either with eyes open or closed, which is why lying or standing with eyes open should not *per se* be interpreted as a sign of a negative state in cattle.

We only included two tail postures in our ethogram since a more detailed recording of the tail was not possible during live observations. As a binary outcome measure, tail posture was possibly not recorded in sufficient detail to capture potential differences between husbandry systems. However, even though the ethogram used by de Oliveira and Keeling (2018) included more categories of tail movements, “tail hanging stationary” was also the most frequent position across all three situations in their study. Movement of the tail in our study was almost only recorded on PASTURE farms, where it was likely displayed to flick away flies (Dougherty et al., 1995).

By analysing co-occurring postures of different body parts with the cspade algorithm, we detected the combinations of postures that were shown most frequently while the animals were inactive. The number of detected combinations including two, three or four body parts was higher for INTENSIVE and SEMI than for PASTURE systems, i.e. cattle on pasture displayed fewer different postures while being inactive than cattle in INTENSIVE and SEMI systems. This result is counter-intuitive at first glance since one would expect animals in more intensive systems to show fewer different postures in line with studies showing that behavioural diversity is reduced in barren compared to enriched systems (e.g. Haskell et al., 1996; Powell, 1995; Wemelsfelder et al., 2000). However, these studies looked at the whole behavioural repertoire, including, if not primarily focusing on, active behaviour, and not on co-occurring postures in inactive animals. Currently we lack information on which combinations of postures are indicative of positive states. It may thus be possible that cattle on PASTURE showed fewer, but potentially positive combinations, for longer compared to cattle in INTENSIVE and SEMI systems, but this explanation is rather speculative and needs further investigation. Consideration of methodological aspects of the experimental design did not help explaining our finding either. Fewer animals and thus fewer data points were considered on PASTURE compared to INTENSIVE and SEMI systems, but the results remained the same when we truncated the data for both number of animals and number of data points and re-ran the analysis. Moreover, the lower variation in combinations shown by cattle on PASTURE may have resulted from certain combinations being more difficult to be identified in this husbandry system, but this explanation is rather unlikely since all single postures were observed on PASTURE.

The six most frequently co-occurring fourfold combinations (i.e. Basic, Head, Ears, Eyes) were shown for approximately three fourth of the inactivity time, while the remaining four fourfold combinations plus the combinations that did not reach the threshold of 0.1 accounted for the remaining 25 % of the time. The pattern was relatively similar for INTENSIVE and SEMI, while it differed for PASTURE, especially with respect to the combination of Lying with the Head up, Ears low and Eyes closed, which was the most frequently observed combination on PASTURE.

Our data give a descriptive overview of what inactivity looks like in fattening cattle with respect to basic body postures, head and ear postures and the closure of the eyes. However, they do not provide information on the relevance of these combinations for cattle welfare. Differences between husbandry systems were mostly subtle, and whether these differences are indicative of positive or negative welfare or whether they are rather characteristics of the respective husbandry systems cannot be answered with our data. Future research should thus specifically investigate the association between positive and negative inactivity and co-occurring postures.

The machine learning algorithm cspade is a promising statistical tool to capture the complexity of behavioural data. With the constant increase in automatically recorded data, for example by various data loggers, an algorithm like cspade may help analysing information simultaneously obtained from different data loggers of the same animal over many time points. In addition to the analysis of co-occurrences of postures of different body parts, it would be valuable to analyse transitions between postures in future studies, since behaviour is not a snap shot in time, but dynamic (Asher et al., 2009). Specifically, we suggest analysing transitions per body part (e.g. transitions between different ear postures) as well as transitions between the simultaneous occurrences of postures from different body parts (e.g. from “Standing with Head up, Ears forwards, Eyes open” to “Lying with Head up, Ears backwards, Eyes open”) to receive a more complete picture of the whole animal and thus to advance our understanding of what different forms of inactive behaviour mean for animal welfare.

## Conclusion

Our study is the first to give an overview of what inactive behaviour in fattening cattle looks like. We conclude that both the principle of the Inactivity Ethogram to look for detailed postures and the machine learning algorithm cspade used to assess co-occurring postures of different body parts are promising tools for future studies on the welfare relevance of inactive behaviour specifically and the identification of indicators of emotional states more generally.

## Acknowledgements

We would like to thank August Bittermann and Julia Trieb for contacting many of the farmers and to all farmers who were willing to participate in our study. Moreover, we would like to thank Christine Leeb for fruitful discussions on the design of the study and Martina Knöbl for the analysis of the videos. We gratefully acknowledge funding received by the Ombuds Office for Animal Protection of the City of Vienna and by the Universities Federation for Animal Welfare (Animal Welfare Student Scholarship awarded to FM).

R scripts and formatted datasheets for the cspade analysis and network figure are available at Newcastle University data repository (https://data.ncl.ac.uk).

